# mRNA vaccines and hybrid immunity use different B cell germlines to neutralize Omicron BA.4 and BA.5

**DOI:** 10.1101/2022.08.04.502828

**Authors:** Emanuele Andreano, Ida Paciello, Giulio Pierleoni, Giuseppe Maccari, Giada Antonelli, Valentina Abbiento, Piero Pileri, Linda Benincasa, Ginevra Giglioli, Giulia Piccini, Concetta De Santi, Claudia Sala, Duccio Medini, Emanuele Montomoli, Piet Maes, Rino Rappuoli

## Abstract

SARS-CoV-2 omicron BA.4 and BA.5, characterized by high transmissibility and ability to escape natural and vaccine induced immunity, are rampaging worldwide. To understand the escape mechanisms, we tested the neutralizing activity against omicron BA.4 and BA.5 of a panel of 482 human monoclonal antibodies that had been isolated from people who received two or three mRNA vaccine doses or from people that had been vaccinated after infection. None of the antibodies isolated after two vaccine doses neutralized omicron BA.4 and BA.5, while these variants were neutralized by approximately 15% of antibodies obtained from people that received three doses or had been vaccinated after infection. Remarkably, the antibodies isolated after three vaccine doses targeted mainly the receptor binding domain (RBD) Class 1/2 epitope region and were encoded by the IGHV1-69 and IGHV3-66 B cell germlines, while the antibodies isolated after infection recognized mostly the RBD Class 3 epitope region and the NTD, and were encoded by the IGHV2-5;IGHJ4-1 and IGHV1-24;IGHJ4-1 germlines. The observation that mRNA vaccination and hybrid immunity elicit a different immunity against the same antigen is intriguing and its understanding may help to design the next generation of therapeutics and vaccines against COVID-19.

## INTRODUCTION

Almost three years after the first case of SARS-CoV-2 in Wuhan in December 2020, 578 million cases and 6.4 million deaths have been reported to be caused by the COVID-19 pandemic. In the meantime, the original Wuhan virus has been replaced by several variants of concern named alpha, beta, gamma, delta and omicron, each characterized by the ability to escape natural and vaccine induced antibody neutralization and by an improved ability to transmit from person to person^1^. Since November 2021 the omicron variant replaced all previous viruses and generated new lineages named BA.1, BA.2, BA.4 and BA.5^1–3^. According to the global initiative on sharing all influenza data (GISAID) database the most recent SARS-CoV-2 omicron sublineages BA.4 and BA.5 are currently the most abundant SARS-CoV-2 circulating variants worldwide^4^. Different scenarios could explain the fast spreading of these new sublineages and the ability to outcompete previous omicron variants. Examples are the lack of the G496S mutation in the Spike (S) protein, which results in increased human angiotensin-converting enzyme 2 (hACE2) binding affinities compared to other omicron variants^5^, the evolution of novel mutations on the S protein, which conferred enhanced resistance to neutralizing antibodies^6^, and the ability to better suppress and antagonize the innate immune defenses^7^. The SARS-CoV-2 neutralizing antibodies target the receptor binding domain (RBD) and N terminal domain (NTD) of the S protein, which is a trimeric glycoprotein exposed on the surface of the virus^8,9^. The SARS-CoV-2 omicron sublineages BA.4 and BA.5 share an identical S glycoprotein which carries 31 mutations on its surface **(Supplementary Fig. 1a)**^6^. As for the initial omicron BA.1 and BA.2, both the receptor binding domain (RBD) and N terminal domain (NTD) immunodominant sites of these new sublineages are heavily mutated. The NTD of BA.4 and BA.5 harbors 9 mutations (29.0%) on this domain which are represented by 4 substitutions (T19I, L24S, G142D and V213G) and 5 deletions (Δ25-27 and Δ69-70). The mutational pattern is extremely similar to the parental BA.2 lineage with the exception of the Δ69-70 mutation which was present in the original BA.1 omicron variant. As observed with all previous SARS-CoV-2 variants of concern, the RBD of omicron BA.4 and BA.5 displays only substituted residues highlighting the more conservative structure of this domain. The RBD carries 17 mutations (54.8%), and 9 of them are within the receptor binding motif (RBM) which spans from residue S438 to Y508 of the S protein^10^ **(Supplementary Fig. 1b)**. Recent studies have described the impact of the omicron variants, including BA.4 and BA.5, on the polyclonal antibody response of subjects infected, vaccinated and with hybrid immunity^11–13^, as well as on a set of 28 and 158 neutralizing antibodies (including therapeutic, database and previous publication-derived antibodies isolated from a variety of subjects and cohorts)^14,15^, or on a library of 1,640 RBD-binding antibodies^5^. In this study we evaluated the neutralizing activity against BA.4 and BA.5 variants of 482 neutralizing human monoclonal antibodies (nAbs) that neutralized the original Wuhan virus. Our data confirm at single cell level that only a minority of nAbs cross-neutralize BA.4 and BA.5 lineages and reveal that the B cell germlines usage and S protein epitopes targeted for cross-neutralization are different in vaccinated and infected people.

## RESULTS

### Neutralization of omicron sublineages

A collection of 482 neutralizing human monoclonal antibodies (nAbs) against the SARS-CoV-2 virus originally isolated in Wuhan, were used in this study. They derived from three different cohorts: SARS-CoV-2 seronegative subjects vaccinated with two (SN2) or three (SN3) doses of COVID-19 mRNA vaccines, and subjects exposed to SARS-CoV-2 infection and subsequently vaccinated with two doses of the same mRNA vaccines (seropositive 2^nd^ dose; SP2)^16,17^. All subjects, with the exception of VAC-010 in the SN3 cohort which was immunized with mRNA-1273, received the BNT162b2 mRNA vaccine^16,17^. Their neutralizing potency against the SARS-CoV-2 omicron variants BA.4 and BA.5, was tested by the cytopathic effect-based microneutralization assay (CPE-MN) against live viruses in biosafety level 3 (BSL3) laboratories. Overall, less than 15% of the antibodies retained a neutralizing activity against the omicron BA.4 and BA.5 variants. As shown in **Fig. 1**, none of the 52 antibodies from the SN2 cohort were able to neutralize omicron BA.4 and BA.5 variants, while minimal cross-protection was observed against BA.1 (*n* = 1; 1.9%) and BA.2 (*n* = 4; 7.7%) **(Fig. 1a)**. Conversely, of the 206 nAbs in the SN3 cohort, 14.6 (*n* = 30) and 14.1% (*n* = 29) cross-neutralized omicron BA.4 and BA.5 respectively **(Fig. 1b-c)**. Similarly, in the case of SP2, 15.5 (*n* = 32) and 14.6% (*n* = 30) of the 224 nAbs cross-neutralized these SARS-CoV-2 omicron variants. The overall nAbs neutralization potency against omicron BA.4 and BA.5 in the SN3 and SP2 groups, reported as geometric mean 100% inhibitory concentration (GM-IC_100_), showed up to 2.62- and 5.34-fold GM-IC_100_ decrease compared to Wuhan and up to 1.62- and 3.02-fold decrease compared to BA.1 and BA.2 respectively **(Fig. 1b-d)**. Interestingly, none of the nAbs tested showed extremely potent neutralization activity (IC_100_ <10 ng ml^-1^) against all omicron viruses.

**Fig. 1.**
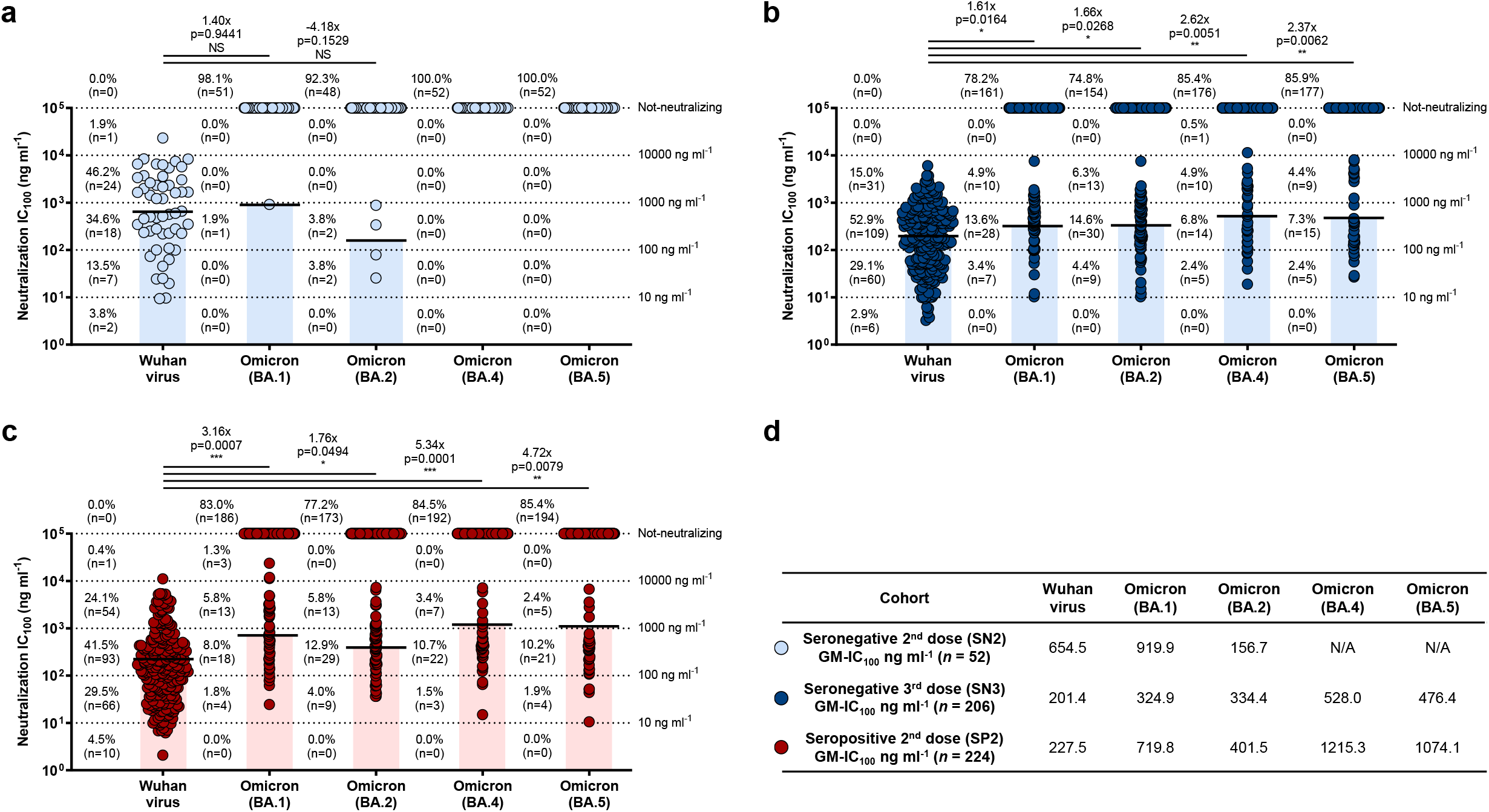
Potency and breadth of neutralization of nAbs against SARS-CoV-2 omicron variants. **a-c**, Scatter dot charts show the neutralization potency, reported as IC_100_ (ng ml^-1^), of nAbs tested against the original Wuhan SARS-CoV-2 virus, and the omicron BA.1, BA.2, BA.4 and BA.5 lineages for SN2 (a), SN3 (b), and SP2 **(c)** respectively. The number, percentage, GM-IC_100_ (black lines and colored bars), fold-change and statistical significance of nAbs are denoted on each graph. Reported fold-change and statistical significance are in comparison with the Wuhan virus. Technical duplicates were performed for each experiment. **d,** The table shows the IC_100_ geometric mean (GM-IC_100_) of all nAbs pulled together from SN2, SN3 and SP2 against all SARS-CoV-2 viruses tested. A nonparametric Mann–Whitney t test was used to evaluate statistical significances between groups. Two-tailed p-value significances are shown as *p < 0.05, **p < 0.01, and ***p <0.001.

### Mapping RBD and NTD cross-protective nAbs

To understand the type of antibodies mainly responsible for cross-protection against the omicron variants, we investigated the neutralization activity of RBD and NTD binding nAbs **(Fig. 2)**. RBD-targeting nAbs were previously classified based on their ability to compete with the Class 1/2 antibody J08^18^, the Class 3 antibody S309^19^, and the Class 4 antibody CR3022^20^, or for their lack of competition with the three tested antibodies (Not-competing)^17,21^. The SN2 group (*n* = 46) showed mainly nAbs targeting the RBD-Class 1/2 epitope region against Wuhan (*n* = 26; 56.5%), BA.1 (*n* = 1; 100%) and BA.2 (*n* = 2; 50%), while no neutralization activity was observed against the BA.4 and BA.5 omicron variants **(Fig. 2a,d)**. A similar trend was also observed for RBD and NTD-targeting nAbs isolated from the SN3 cohort (*n* = 197). Indeed, RBD-Class 1/2 targeting nAbs represented the 49.7% (*n* = 98) of antibodies neutralizing the Wuhan virus, and their percentage increased against the omicron sublineages constituting the 62.8 (*n* = 27), 52.0 (*n* = 26), 58.3 (*n* = 14) and 56.5% (*n* = 13) of nAbs able to cross-neutralize the omicron BA.1, BA.2, BA.4 and BA.5 respectively **(Fig. 2b,e)**. Interestingly, while NTD-targeting antibodies were the second most abundant class among Wuhan nAbs, they lost almost completely their functionality against the omicron lineages, representing only 2.3 (*n* = 1) and 4.0% (*n* = 2) of nAbs against BA.1 and BA.2, while showing no activity towards the omicron BA.4 and BA.5 **(Fig. 2b,e)**. Interestingly, RBD and NTD-targeting nAbs isolated from the SP2 cohort (*n* = 215) showed a completely different profile against the omicron variants. In fact, cross-neutralizing antibodies targeted preferentially the RBD-Class 3 epitope region and constituted the 51.0 (*n* = 26), 56.3 (*n* = 18) and 56.7% (*n* = 17) of nAbs able to neutralize omicron BA.2, BA.4 and BA.5 respectively **(Fig. 2c,f)**. In addition, differently from what observed in the SN3 cohort, NTD-targeting nAbs isolated in the SP2 group retained high level of functionality against the omicron BA.2, BA.4 and BA.5, representing the 15.7 (*n* = 8), 28.1 (*n* = 9) and 26.7% (*n* = 8) cross-protective nAbs against these variants respectively **(Fig. 2c,f)**. The RBD-Class 1/2 antibodies that were the most abundant in neutralizing Wuhan (*n* = 114; 53.0%) and BA.1 (*n* = 19; 50.0%), were heavily escaped by omicron BA.2, BA.4 and BA.5, representing only the 31.4 (*n* = 16), 15.6 (*n* = 5) and 16.7% (*n* = 5) of nAbs respectively **(Fig. 2c,f)**. Finally, independently from their overall frequency, the neutralization potency of omicron BA.4 and BA.5 of Class 1/2 and Class 3 nAbs in the SN3 group was higher than in the SP2 group **(Supplementary Fig. 2)**. Noteworthy, while no NTD-targeting antibodies able to neutralize BA.4 and BA.5 were found in the SN3 group, nAbs isolated from the SP2 cohort that targeted this S protein domain were the second most abundant group of antibodies and showed a neutralization potency comparable to Class 3 nAbs and up to 2.44-fold higher GM-IC_100_ compared to Class 1/2 antibodies isolated in this cohort **(Supplementary Fig. 2)**.

**Fig. 2.**
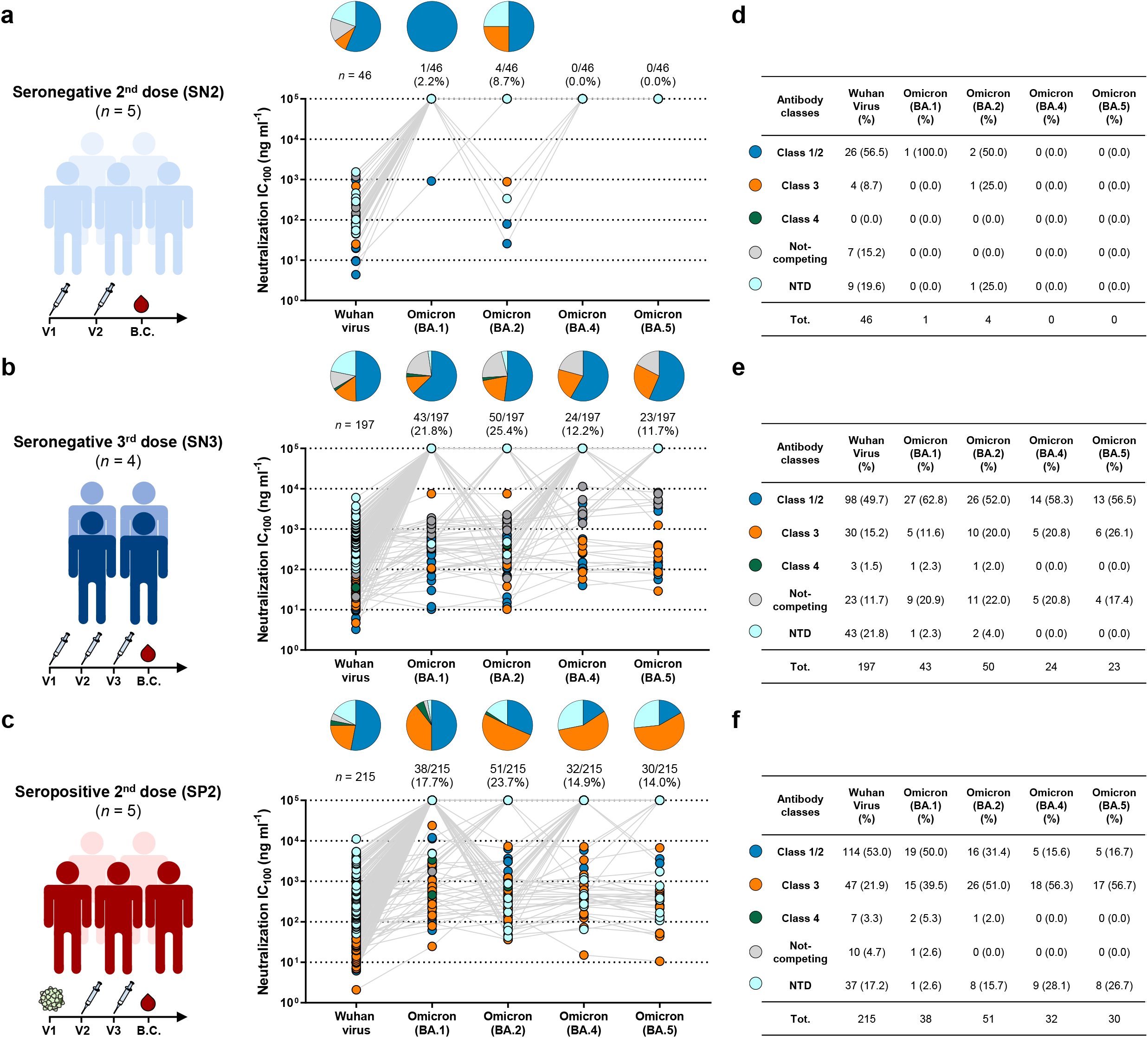
Distribution of RBD and NTD-targeting nAbs against omicron variants. **a-c**, Pie charts show the distribution of cross-protective nAbs based on their ability to bind Class 1/2 (blue), Class 3 (orange) and Class 4 (dark green) regions on the RBD, as well as not-competing nAbs (gray) and NTD-targeting nAbs (cyan). Dot charts show the neutralization potency, reported as IC_100_ (ng m^-1^), of nAbs against the Wuhan virus and the omicron BA.l, BA.2, BA.4 and BA.5 variants observed in the SN2 (a), SN3 (b) and SP2 (c) cohorts. The number and percentage of nAbs are denoted on each graph. **d-f,** tables summarize number and percentage of Class 1/2, Class 3, Class 4, not-competing and NTD-targeting nAbs for each tested variant in the SN2 (d), SN3 (e) and SP2 (f) cohorts.

### B cell germline usage of cross-neutralizing omicron nAbs

In addition to the functional characterization and epitope mapping analyses, we investigated the B cell germline and V-J gene rearrangements (IGHV;IGHJ) used by highly cross-reactive nAbs against omicron variants. Of the 482 nAbs assessed in this study we previously recovered 430 heavy chain sequences: 46 from SN2, 176 from SN3 and 208 from SP2 ^16,17^. In SN2 subjects, predominant B cell germlines neutralizing the Wuhan strain include IGHV1-69;IGHJ4-1, IGHV3-30;IGHJ6-1, IGHV3-53;IGHJ6-1, IGHV3-66;IGHJ4-1 B cell germlines^16,22–25^. These germlines constituted the 28.3% of nAbs able to neutralize the Wuhan virus and all of them, lost completely their functional activity against all omicron variants; the only exception was one nAb encoded by the IGHV3-53;IGHJ6-1 germline which was able to neutralize with high potency BA.2 **(Fig. 3a; Supplementary Table 1; Supplementary Table 2)**. Differently from the SN2 group, the SN3 and SP2 cohorts contained SARS-CoV-2 cross-neutralizing nAbs against all omicron variants. In the SN3 cohort, omicron cross-neutralizing antibodies were dominated by five V-J gene rearrangements. These were IGHV1-58;IGHJ3-1, IGHV1-69;IGHJ3-1, IGHV1-69;IGHJ4-1, IGHV3-66;IGHJ4-1 and IGHV3-66;IGHJ6-1 **(Fig. 3b; Supplementary Table 1**). These five germlines are well known and encode for potently neutralizing RBD-targeting Class 1 and Class 2 nAbs ^20,21,23,26^. These germlines represented the 32.4% of nAbs against the original Wuhan virus, and the 58.3, 54.8, 54.6 and 54.6% of cross-neutralizing nAbs against omicron BA.1, BA.2, BA.4 and BA.5 respectively **(Fig. 3b; Supplementary Table 1)**. When we analyzed the neutralization potency of these abundant germlines we observed that the IGHV1-69;IGHJ4-1 was the most abundant germline among all omicron variants, while nAbs encoded by the IGHV3-66;IGHJ6-1 V-J genes were the only to maintain a GM-IC_100_ against all omicron variants similar to what observed for the original Wuhan virus **(Fig. 3b; Supplementary Table 2)**. The remaining germlines showed a 1.67- to 45.43-fold reduction in their GM-IC_100_ compared to the Wuhan virus. For the SP2 cohort, omicron cross-functional antibodies derived mainly from three germlines which differed from those found in the SN3 group. These germlines used the IGHV1-24;IGHJ6-1, IGHV1-58;IGHJ3-1 and IGHV2-5;IGHJ4-1 V-J gene rearrangements. nAbs encoded by these B cell germlines represent only 11.5% of all antibodies against the Wuhan virus **(Fig. 3c; Supplementary Table** 1) and their frequency increased to 24.3, 28.6, 30.0 and 31.0% for cross-neutralizing nAbs against omicron BA.1, BA.2, BA.4 and BA.5 respectively. The IGHV1-58;IGHJ3-1 and IGHV2-5;IGHJ4-1 germlines encoded for RBD-targeting Class 1 and Class 3 nAbs respectively^16,21,26^, while the IGHV1-24;IGHJ6-1 V-J gene rearrangement is mainly used by NTD-targeting antibodies^27,28^. With the exception of BA.1, the IGHV2-5;IGHJ4-1 germline is the most frequently used by omicron cross-neutralizing nAbs isolated in this cohort. In addition, antibodies carrying the IGHV2-5;IGHJ4-1 rearrangement showed to be the only group of nAbs, among the three highly frequent germlines in the SP2 cohort, able to cross-neutralize all omicron variants although showing a 7.04 – 13.51-fold decrease in GM-IC_100_ compared to the Wuhan virus **(Supplementary Table 1; Supplementary Table 2)**.

**Fig. 3.**
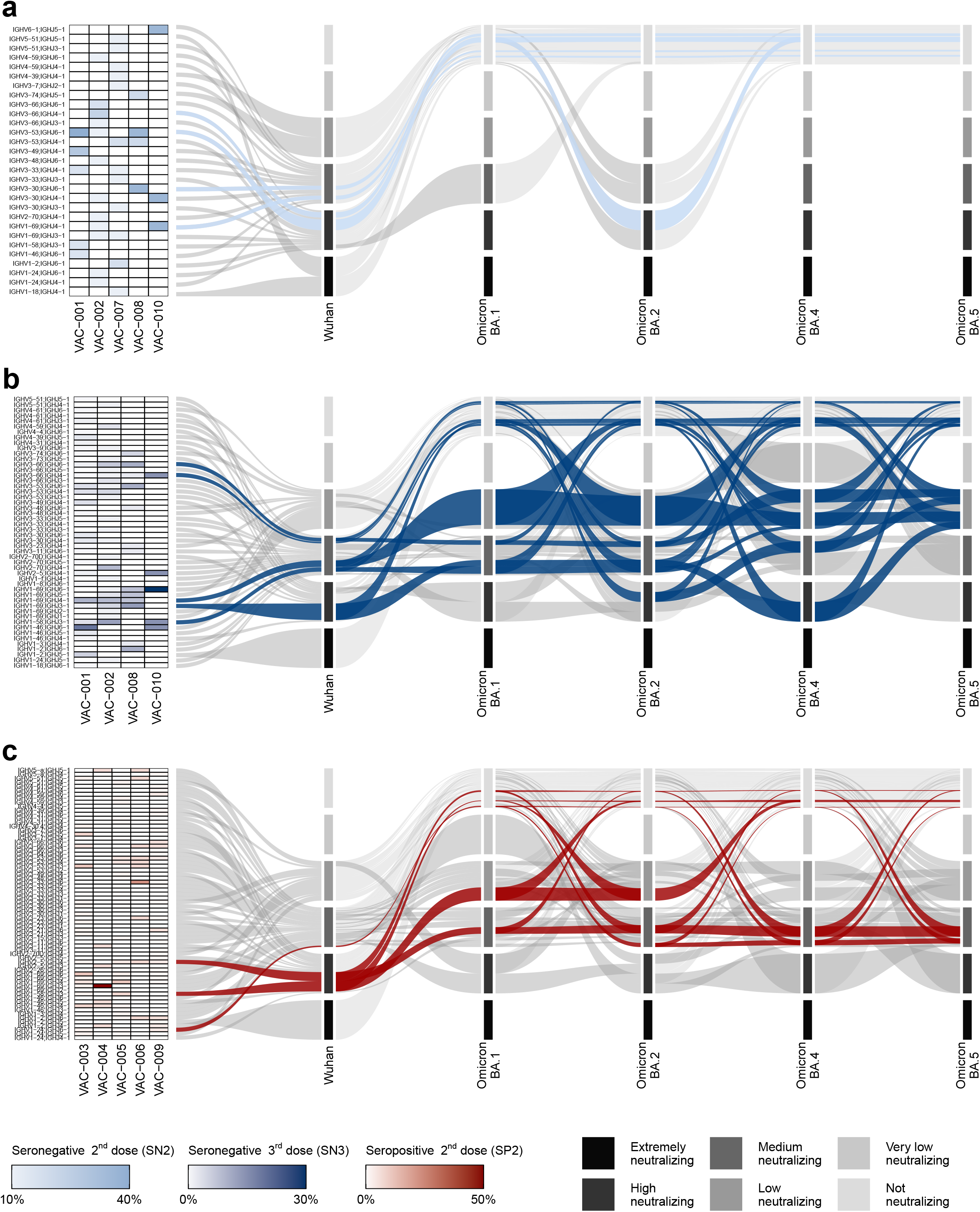
IGHV;IGHJ gene usage of omicron cross-neutralizing antibodies. **a-c**, Heatmaps and alluvial plots display the antibody IGHV;IGHJ gene rearrangements frequency for each single donors and for pulled nAbs respectively for SN2 (a), SN3 (b) and SP2 (c). In the alluvial plots, V-J gene rearrangements were highlighted if they represented at least 10% of all antibodies able to cross-neutralize a specific omicron variant. Below this threshold, several germlines showed identical or similar frequency values and therefore were not considered as predominant. Selected germlines were highlighted as light blue, dark blue and red for SN2, SN3 and SP2 respectively. Germline usage is shown for nAbs against the original Wuhan virus and omicron BA.1, BA.2, BA.4 and BA.5 variants. For each variant, nAbs were grouped into six different categories (strata) based on the neutralization potency (GM-IC_100_) of all nAbs encoded by the specific germline. Strata are defined as extremely neutralizing (≤ 10 ng ml-1), high neutralizing (≤ 100 ng ml-1), medium neutralizing (≤ 1,000 ng ml-1), low neutralizing (≤ 10,000 ng ml-1), very low neutralizing (< 100,000 ng ml-1) and not neutralizing (≥ 100,000 ng ml-1). The flow size indicates the frequency of the specific germline within the strata to which it is linked.

### Impact on therapeutic nAbs

Since the RBD and RBM are heavily mutated in the omicron BA.4 and BA.5 variants, and they represent the major targets of antibodies approved for clinical treatment of COVID-19, we evaluated the impact of omicron BA.4 and BA.5 mutations on eight therapeutic mAbs approved for therapy. Specifically, we tested three Class 1 mAbs, REGN10933 (Casirivimab)^29^, LY-CoV016 (Etesevimab)^30^, and COV2-2196 (Tixagevimab)^31^, the Class 2 targeting nAb LY-CoV555 (Bamlanivimab)^32^, and four Class 3 directed nAbs, S309 (Sotrovimab)^19^, REGN10987 (Imdevimab)^29^, LY-CoV1404 (Bebtelovimab)^33^ and COV2-2130 (Cilgavimab)^31^, by CPE-MN against the live SARS-CoV-2 virus originated in Wuhan and the omicron viruses **(Supplementary Fig. 3)**. All tested nAbs showed neutralization activity against the ancestral Wuhan virus with a 100% inhibitory concentration (IC_100_) ranging from 19.5 to 176.8 ng ml^-1^. Class 1 and Class 2 nAbs, derived from the IGHV3-11;IGHJ4-1 (Casirivimab), IGHV3-66;IGHJ4-1 (Etesevimab), IGHV1-58;IGHJ3-1 (Tixagevimab), and IGHV1-69;IGHJ6-1 (Bamlanivimab) germlines, were evaded by all omicron variants. Differently, Class 3 antibodies, encoded by the IGHV1-18;IGHJ4-1 (Sotrovimab), IGHV3-30;IGHJ4-1 (Imdevimab), IGHV2-5;IGHJ1-1 (Bebtelovimab), and IGHV3-15;IGHJ4-1 (Cligavimab) germlines, retained their neutralization activity against at least one omicron variant. Indeed, S309 (Sotrovimab) was able to neutralize the omicron BA.1 virus with a 3.17-fold reduction, while no activity was detected against the other omicron variants. REGN10987 (Imdevimab) and COV2-2130 (Cilgavimab), neutralized three out of four variants despite showing up to 81.31- and 5.65-fold decrease in their respective IC_100_. Finally, LY-CoV1404 (Bebtelovimab), was the only antibody with high neutralization potency against all omicron lineages showing an IC_100_ of 11.1, 15.6, 44.2 and 62.5 ng ml^-1^ against omicron BA.1, BA.2, BA.4 and BA.5 respectively **(Supplementary Fig. 3)**. These results are in line with previously published works^5,14,15,33^.

## DISCUSSION

In this work, we took advantage of our unique panel of 482 SARS-CoV-2 neutralizing human monoclonal antibodies to address at single cell level the cross-neutralizing properties against the omicron variants of B cells induced by vaccine or hybrid immunity. Our nAb panel, built during the last three years of the COVID-19 pandemic, were identified from people receiving two or three mRNA vaccine doses, and from SARS-CoV-2 infected people that had been subsequently vaccinated with the BNT162b2 mRNA vaccine^16,17^. In agreement with previous studies performed mainly with whole sera^6–8^, we observed that two mRNA vaccine doses were not sufficient to mount a protective antibody response against omicron variants. Conversely, three mRNA vaccine doses and hybrid immunity showed to induce similar, although limited, levels of protection against the omicron variants, with an overall average of 18.9 and 17.5% of nAbs still able to neutralize these viruses for SN3 and SP2 respectively. The observation that vaccination and hybrid immunity show similar level of omicron cross-protection is not aligned with previous studies which, through the analyses of the polyclonal response of subjects with heterologous history of vaccination and infection, reported higher protection in this latter cohort^2,6,9^. Despite the similarity in the number of antibodies neutralizing the omicron variants BA.4 and BA.5 in SN3 and SP2, our analyses revealed dramatic differences in the antibody and B cell germline profiles behind their respective responses. Three mRNA vaccine doses expanded mainly RBD-targeting Class 1/2 nAbs and showed a more clonal B cell response constituted mainly by five germlines (IGHV1-58;IGHJ3-1, IGHV1-69;IGHJ3-1, IGHV1-69;IGHJ4-1, IGHV3-66;IGHJ4-1 and IGHV3-66;IGHJ6-1) which represented almost 60% of all omicron cross-neutralizing antibodies. Interestingly, two of the five abundant germlines encoding for cross-neutralizing nAbs in SN3 (IGHV1-69;IGHJ4-1, IGHV3-66;IGHJ4-1) were also present in the SN2 cohort where they showed no functional activity against omicron variants. This suggests that a third mRNA vaccine dose enhances B cell affinity maturation of selected germlines and drive their expansion and subsequent production of cross-protective nAbs. As for hybrid immunity, RBD-directed Class 3 nAbs and NTD-targeting nAbs were preferentially used and a more diversified B cell response was observed. In fact, only three major germlines were identified (IGHV1-24;IGHJ6-1, IGHV1-58;IGHJ3-1 and IGHV2-5;IGHJ4-1) which represented no more than 31% of the whole antibody response against the omicron variants. The observation that homologous mRNA vaccination and infection drive the expansion of different B cell germlines which produce nAbs targeting distinct epitopes on the SARS-CoV-2 S protein raises interesting questions about the mechanistic of antigen presentation. Indeed, in both cases the antigen is produced by the host cells and its presentation to the immune cells is likely to derive from the different cell types expressing the antigen, the stabilization of the S protein in its prefusion conformation following the insertion of two prolines, the absence of other viral components and the inflammatory environment present during infection. Overall, our work provides unique information on the B cell and antibody response induced by vaccine and hybrid immunity, highlighting similarities and key differences between these two immunologically distinct cohorts that could be exploited for the design of next generation therapeutics and vaccines against SARS-CoV-2.

## METHODS

### Enrollment of COVID-19 vaccinees and human sample collection

Human samples from COVID-19 infected and vaccinated donors, who received two or three vaccine doses, of both sexes, who gave their written consent, were previously collected through a collaboration with the Azienda Ospedaliera Universitaria Senese, Siena (IT)^16,17^. Subjects in the seropositive 2^nd^ dose cohort resulted positive to SARS-CoV-2 infection between October and November 2020^16^. The study was approved by the Comitato Etico di Area Vasta Sud Est (CEAVSE) ethics committees (Parere 17065 in Siena) and conducted according to good clinical practice in accordance with the declaration of Helsinki (European Council 2001, US Code of Federal Regulations, ICH 1997). This study was unblinded and not randomized. No statistical methods were used to predetermine sample size.

### SARS-CoV-2 authentic viruses neutralization assay

All SARS-CoV-2 authentic virus neutralization assays were performed in the biosafety level 3 (BSL3) laboratories at Toscana Life Sciences in Siena (Italy) and Vismederi Srl, Siena (Italy), which are approved by a Certified Biosafety Professional and inspected annually by local authorities. To assess the neutralization potency and breadth of nAbs against the live SARS-CoV-2 viruses, a cytopathic effect-based microneutralization assay (CPE-MN) was performed as previously described^16,17,21,22^. Briefly, nAbs were co-incubated with SARS-CoV-2 viruses used at 100 median Tissue Culture Infectious Dose (100 TCID_50_) for 1 hour at 37°C, 5% CO_2_. The mixture was then added to the wells of a 96-well plate containing a sub-confluent Vero E6 cell monolayer. Plates were incubated for 3-4 days at 37°C in a humidified environment with 5% CO_2_, then examined for CPE by means of an inverted optical microscope by two independent operators. TAP expressed nAbs were tested at a starting dilution of 1:5 and diluted step 1:2. Flask expressed nAbs were tested at a starting concentration of 2 μg m^-1^ and diluted step 1:2. Technical duplicates and triplicates were performed to evaluate the IC_100_ of TAP and purified nAbs respectively. In each plate positive and negative control were used as previously described^16,17,21,22^.

### SARS-CoV-2 virus variants CPE-MN neutralization assay

The SARS-CoV-2 viruses used to perform the CPE-MN neutralization assay were the Wuhan (SARS-CoV-2/INMI1-Isolate/2020/Italy: MT066156), omicron BA.1 (GISAID ID: EPI_ISL_6794907), BA.2 (GISAID ID: EPI_ISL_10654979), BA.4 (GISAID ID: EPI_ISL_13360709) and BA.5 (GISAID ID: EPI_ISL_13389618).

### Single cell RT-PCR and Ig gene amplification and transcriptionally active PCR expression

Previously obtained PCRII products^16,17^ were used to recover the antibody heavy and light chain sequences, through Sanger sequencing, and for antibody transcriptionally active PCR (TAP) expression into recombinant IgG1^34^. TAP reaction was performed using 5 μL of Q5 polymerase (NEB), 5 μL of GC Enhancer (NEB), 5 μL of 5X buffer,10 mM dNTPs, 0.125 μL of forward/reverse primers and 3 μL of ligation product, using the following cycles: 98°/2’, 35 cycles 98°/10”, 61°/20”, 72°/1’ and 72°/5’. TAP products were purified and subsequently quantified by Qubit Fluorometric Quantitation assay (Invitrogen). Transient transfection was performed using Expi293F cell line (Thermo Fisher) following manufacturing instructions.

### Flask expression and purification of human monoclonal antibodies

Plasmids carrying the antibody heavy and light chain of nAbs were used for transient transfection of Expi293F™ cells (Thermo Fisher) as previously described^22^. Briefly, cells were grown for six days at 37 °C with 8% CO_2_ shaking at 125 rpm according to the manufacturer’s protocol. Six days after transfection, cell cultures were harvested and clarified by centrifugation (1,110 rpm for 8 min at RT). Cell supernatants were recovered, filtered with 0.45 μm filters to remove particulate material, and then purified through affinity chromatography using a 1 mL HiTrap Protein G HP column (GE Healthcare Life Sciences). Antibodies were eluted from the column using 0.1 M glycine-HCl, pH 2.7. Protein-containing fractions were pooled and dialyzed in PBS buffer overnight at 4°C. Final antibody concentrations were determined by measuring the A562 using Pierce™ BCA Protein Assay Kit (Thermo Scientific). Purified antibodies were stored at −80°C prior to use.

### Functional repertoire analyses

nAbs VH and VL sequence reads were manually curated and retrieved using CLC sequence viewer (Qiagen). Aberrant sequences were removed from the data set. Analyzed reads were saved in FASTA format and the repertoire analyses was performed using Cloanalyst (http://www.bu.edu/computationalimmunology/research/software/)^35,36^.

### Alluvial plot of germline neutralization potency distribution

Alluvial plots were generated to display the neutralization potency distribution of IGHV;IGHJ germlines among the three analyzed cohorts: seronegative 2^nd^ dose (SN2), 3^rd^ dose (SN3) and seropositive 2^nd^ dose (SP2). For each variant indicated on the ordinates, six different categories (strata) are represented to group antibody neutralization potency depending on the average germline IC_100_: extremely neutralizing (≤ 10 ng ml^-1^), high neutralizing (≤ 100 ng ml^-1^), medium neutralizing (≤ 1,000 ng ml^-1^), low neutralizing (≤ 10,000 ng ml^-1^), very low neutralizing (< 100,000 ng ml^-1^) and not neutralizing (≥ 100,000 ng ml^-1^). The germline frequency for each single strata for each variant is represented by the flow size. For the functional antibody repertoire analyses of these two groups, we highlighted the V-J gene rearrangements of nAbs representing at least 10% of all antibodies able to cross-neutralize a specific omicron variant. Below this threshold, several germlines showed identical or similar frequency values and therefore, were not considered as predominant. Selected germlines were colored in light blue, dark blue and red for SN2, SN3 and SP2 respectively. The heatmap on the left represents germline frequency for each individual subject. The figure was assembled with ggplot2 v3.3.5.

### Statistical analysis

Statistical analysis was assessed with GraphPad Prism Version 8.0.2 (GraphPad Software, Inc., San Diego, CA). Nonparametric Mann-Whitney t test was used to evaluate statistical significance between the two groups analyzed in this study. Statistical significance was shown as * for values ≤ 0.05, ** for values ≤ 0.01, and *** for values ≤ 0.001.

## Acknowledgments

This work received funding by the European Research Council (ERC) advanced grant (agreement number 787552 (vAMRes)), and the Italian Ministry of Health (COVID-2020-12371817 project). In addition, this work was supported by a fundraising activity promoted by Unicoop Firenze, Coop Alleanza 3.0, Unicoop Tirreno, Coop Centro Italia, Coop Reno e Coop Amiatina. Piet Maes acknowledges support from the Research Foundation Flanders (COVID19 research grant G0H4420N) and Internal Funds KU Leuven (grant 3M170314).

## Author contributions

Conceived the study: E.A. and R.R.; TAP and flask expression of monoclonal antibodies: I.P., G.A. and V.A.; Recovered VH and VL sequences and performed the repertoire analyses: P.P., E.A. and G.M.; Performed neutralization assays in BSL3 facilities: E.A., I.P., G. Pie., G.A., G.Pic., L.B. and G.G.; Supported day-by-day laboratory activities and management: C.D.S.; Manuscript writing: E.A. and R.R.; Final revision of the manuscript: E.A., I.P., G.Pie., G.M., G.A., V.A., P.P., L.B., G.G., G.Pic., C.D.S., C.S., D.M., E.M., P.M., and R.R.; Coordinated the project: E.A., D.M., E.M., P.M. and R.R.

## Additional information

**Correspondence and requests for materials** should be addressed to R.R.

## Data availability

Source data are provided with this paper. All data supporting the findings in this study are available within the article or can be obtained from the corresponding author upon request.

## SUPPLEMENTARY FIGURES

**Supplementary Fig. 1.**
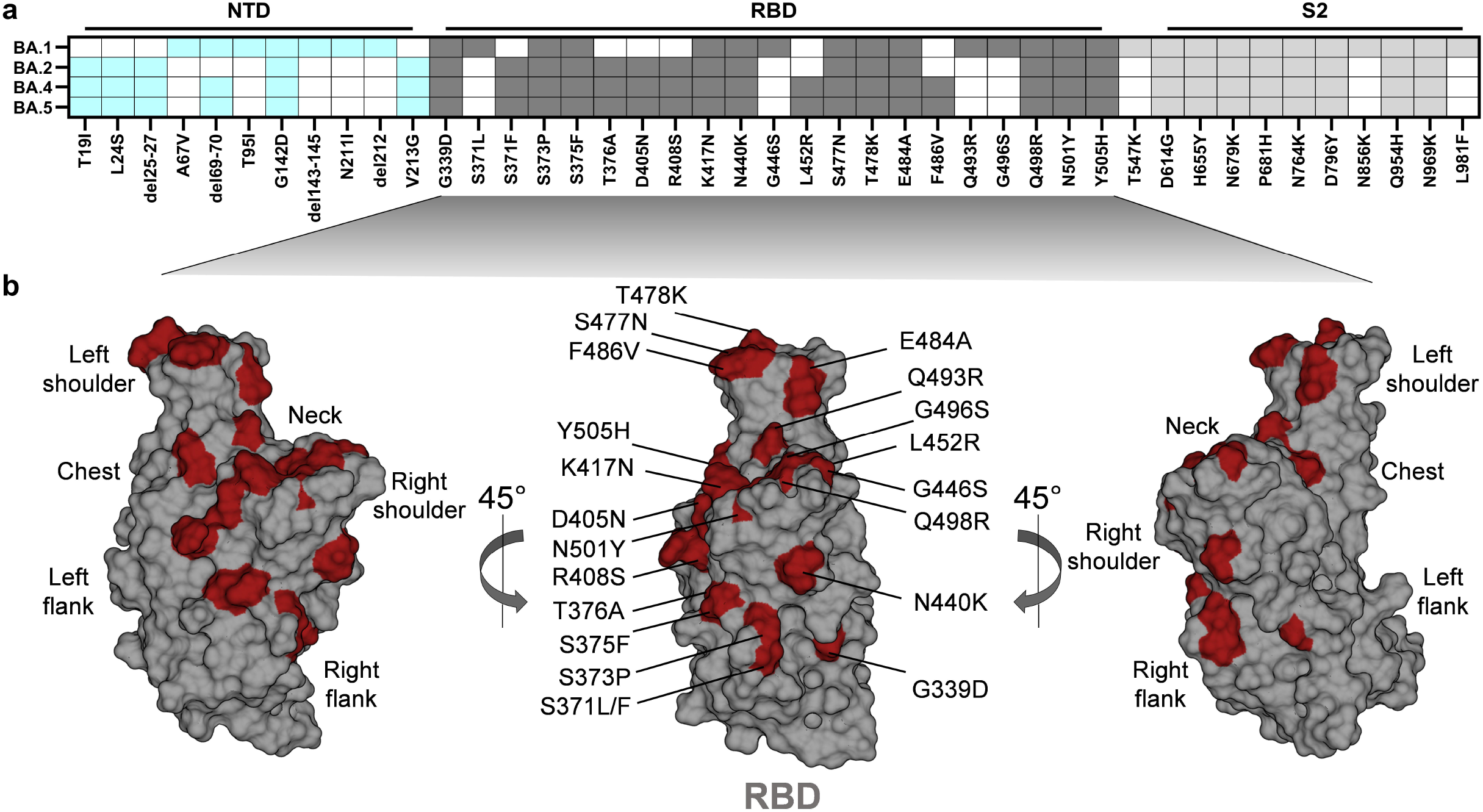
Distribution of omicron mutations on SARS-CoV-2 S protein. **a**, the heatmap shows the SARS-CoV-2 S protein mutations in the NTD (cyan), RBD (dark gray) and S2 domain (light gray) of omicron BA.1, BA.2, BA.4 and BA.5. **b**, Representation of the RBD with all omicron mutations highlighted in red.

**Supplementary Fig. 2.**
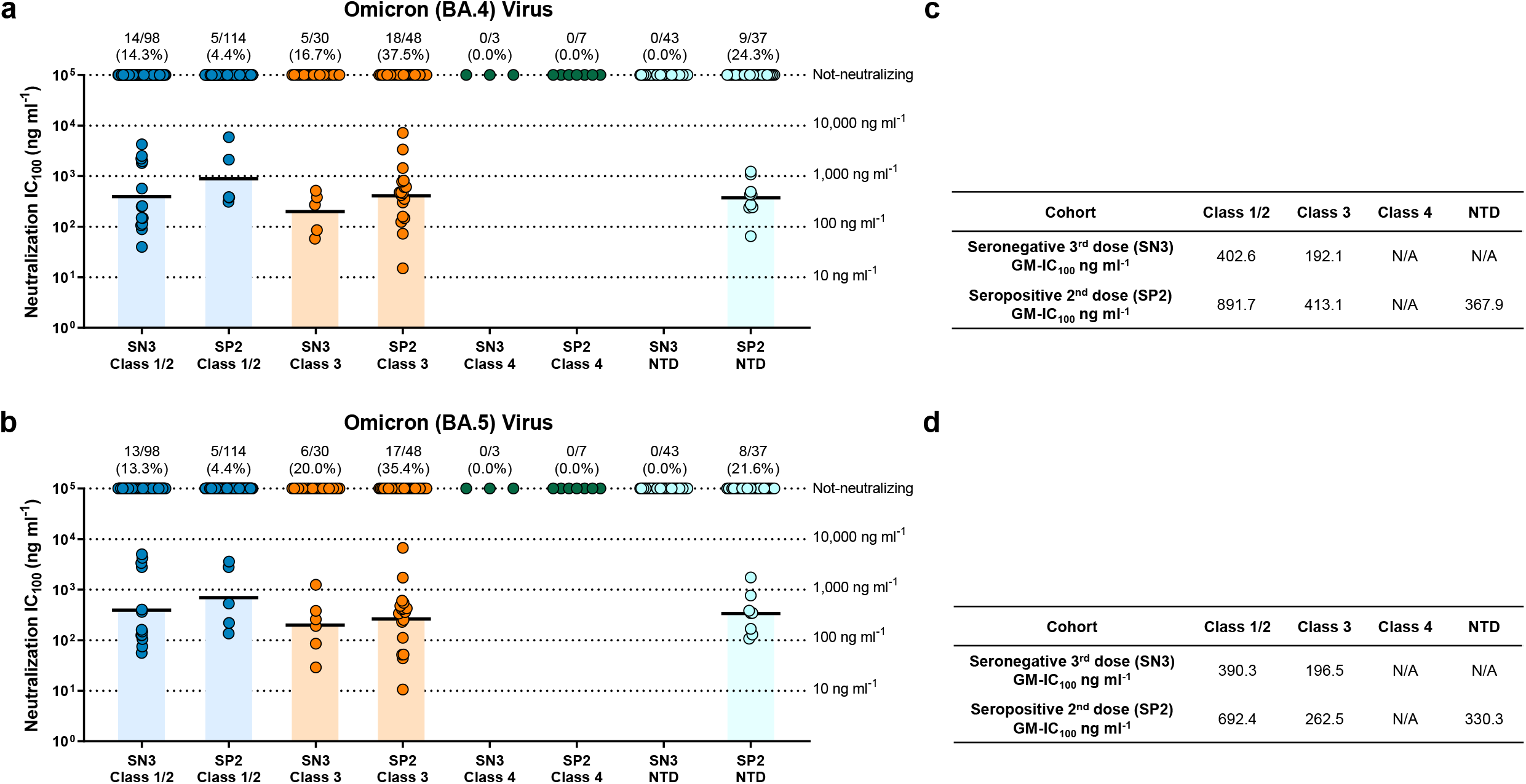
Distribution and neutralization potency of RBD and NTD-targeting nAbs against omicron BA.4 and BA.5. **a-b**, dot charts compare the distribution of nAbs isolated from subjects belonging to the SN3 and SP2 cohorts against omicron BA.4 **(a)** and BA.5 **(b)** omicron variants. The number, percentage and GM-IC_100_ (black lines and colored bars) of nAbs are denoted on each graph. **c-d**, table summarizing the GM-IC_100_ of Class 1/2, Class 3, Class 4 and NTD nAbs against BA.4 (c) and BA.5 (d).

**Supplementary Fig. 3.**
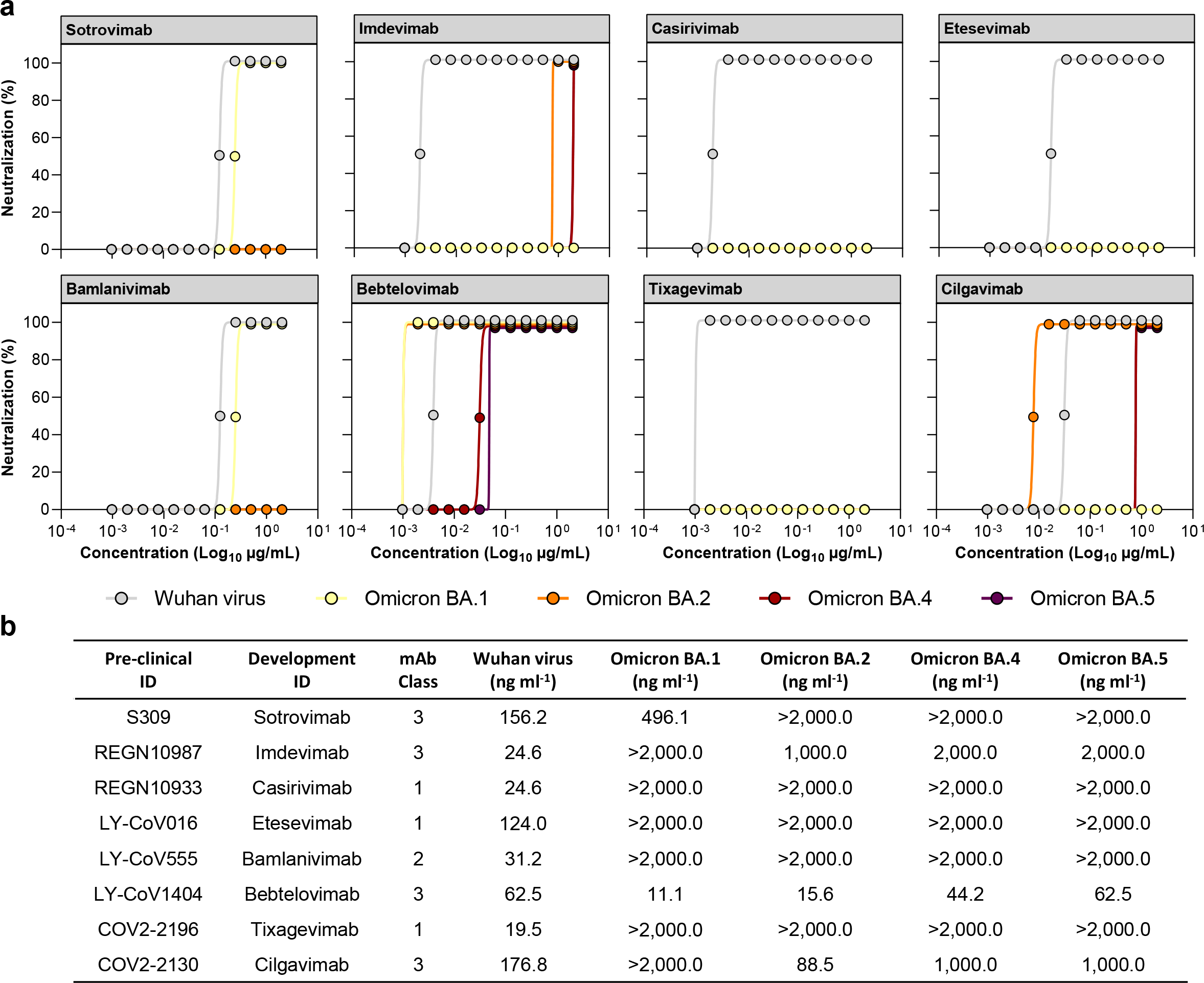
Neutralization activity of COVID-19 therapeutic nAbs. **a**, Graphs show the CPE-MN neutralization activity of therapeutic nAbs against the original SARS-CoV-2 virus originated in Wuhan and the omicron BA.1, BA.2, BA.4 and BA.5 variants. Technical triplicates were performed for each experiment. **b**, the table summarizes the neutralization potency of tested nAbs reported as IC_100_ ng ml^-1^.

## SUPPLEMENTARY TABLES

**Supplementary Table 1.**
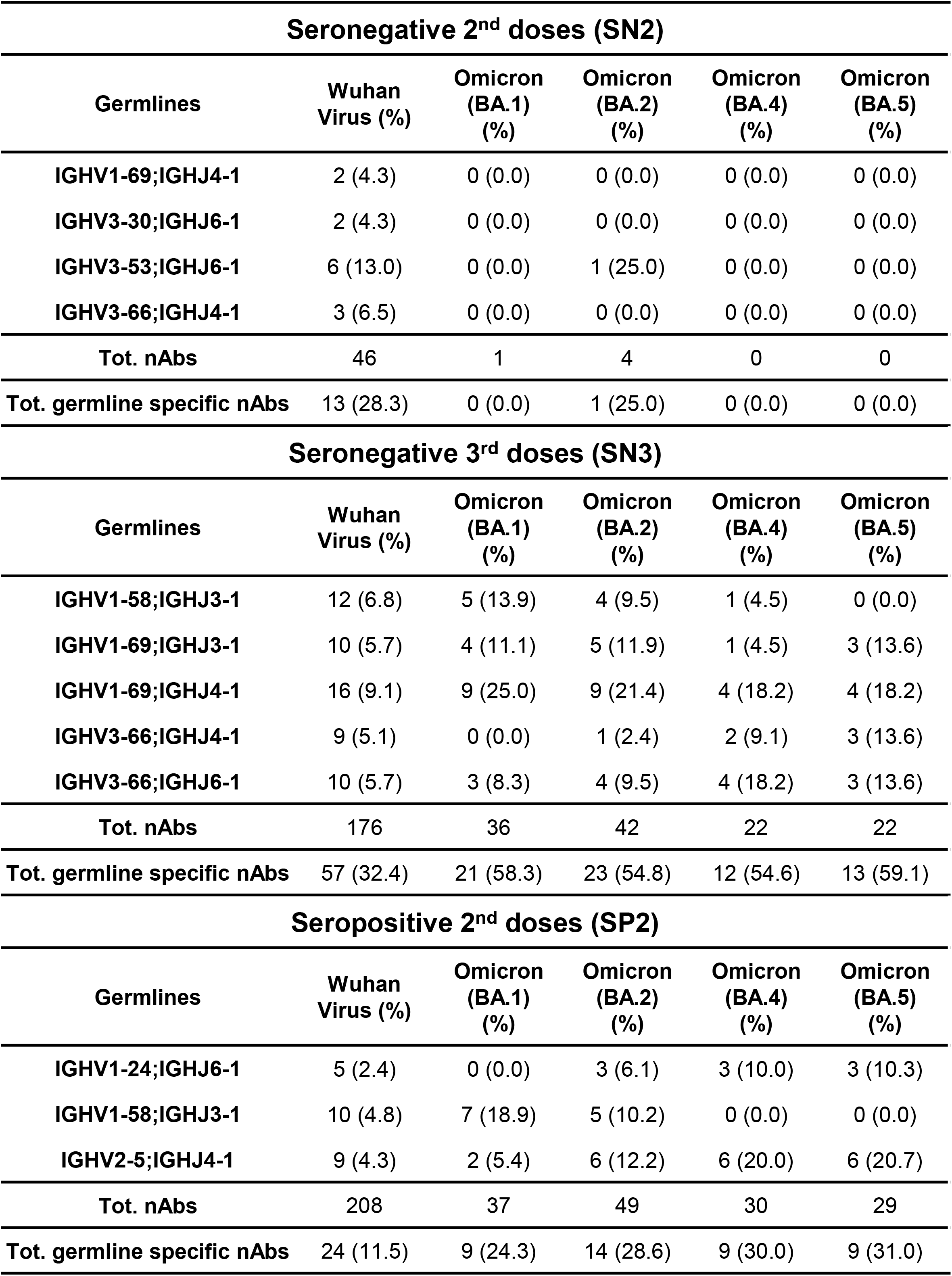
Frequency of predominant germlines against omicron variants.

**Supplementary Table 2.** Neutralization potency of predominant germlines against omicron variants.

Seronegative 2<sup>nd</sup> doses (SN2)
| Germlines | Wuhan<br>Virus<br>GM-IC<sub>100</sub> ng ml<sup>-1</sup> | Omicron<br>(BA.1)<br>GM-IC<sub>100</sub> ng ml<sup>-1</sup> | Omicron<br>(BA.2)<br>GM-IC<sub>100</sub> ng ml<sup>-1</sup> | Omicron<br>(BA.4)<br>GM-IC<sub>100</sub> ng ml<sup>-1</sup> | Omicron<br>(BA.5)<br>GM-IC<sub>100</sub> ng ml<sup>-1</sup> |
| --- | --- | --- | --- | --- | --- |
| IGHV1-69;IGHJ4-1 | 191.63 | N/A | N/A | N/A | N/A |
| IGHV3-30;IGHJ6-1 | 382.71 | N/A | N/A | N/A | N/A |
| IGHV3-53;IGHJ6-1 | 79.58 | N/A | 25.60 | N/A | N/A |
| IGHV3-66;IGHJ4-1 | 66.42 | N/A | N/A | N/A | N/A |

Seronegative 3<sup>rd</sup> doses (SN3)
| Germlines | Wuhan<br>Virus<br>GM-IC<sub>100</sub> ng ml<sup>-1</sup> | Omicron<br>(BA.1)<br>GM-IC<sub>100</sub> ng ml<sup>-1</sup> | Omicron<br>(BA.2)<br>GM-IC<sub>100</sub> ng ml<sup>-1</sup> | Omicron<br>(BA.4)<br>GM-IC<sub>100</sub> ng ml<sup>-1</sup> | Omicron<br>(BA.5)<br>GM-IC<sub>100</sub> ng ml<sup>-1</sup> |
| --- | --- | --- | --- | --- | --- |
| IGHV1-58;IGHJ3-1 | 93.79 | 157.36 | 192.01 | 4261.36 | N/A |
| IGHV1-69;IGHJ3-1 | 50.21 | 155.79 | 124.13 | 39.67 | 616.76 |
| IGHV1-69;IGHJ4-1 | 212.38 | 987.84 | 1069.88 | 1318.53 | 1927.94 |
| IGHV3-66;IGHJ4-1 | 327.61 | N/A | 66.17 | 253.58 | 578.35 |
| IGHV3-66;IGHJ6-1 | 272.30 | 295.43 | 223.63 | 494.97 | 453.70 |

Seropositive 2<sup>nd</sup> doses (SP2)
| Germlines | Wuhan<br>Virus<br>GM-IC<sub>100</sub> ng ml<sup>-1</sup> | Omicron<br>(BA.1)<br>GM-IC<sub>100</sub> ng ml<sup>-1</sup> | Omicron<br>(BA.2)<br>GM-IC<sub>100</sub> ng ml<sup>-1</sup> | Omicron<br>(BA.4)<br>GM-IC<sub>100</sub> ng ml<sup>-1</sup> | Omicron<br>(BA.5)<br>GM-IC<sub>100</sub> ng ml<sup>-1</sup> |
| --- | --- | --- | --- | --- | --- |
| IGHV1-24;IGHJ6-1 | 314.04 | N/A | 130.20 | 413.36 | 463.99 |
| IGHV1-58;IGHJ3-1 | 52.19 | 852.49 | 982.36 | N/A | N/A |
| IGHV2-5;IGHJ4-1 | 28.10 | 379.68 | 197.85 | 264.09 | 166.37 |

